# PARAMO pipeline: reconstructing ancestral anatomies using ontologies and stochastic mapping

**DOI:** 10.1101/553370

**Authors:** Sergei Tarasov, István Mikó, Matthew Jon Yoder, Josef C. Uyeda

## Abstract

Comparative phylogenetics has been largely lacking a method for reconstructing the evolution of phenotypic entities that consist of ensembles of multiple discrete traits – entire organismal anatomies or organismal body regions. In this study, we provide a new approach named *PARAMO* (***P**hylogenetic **A**ncestral **R**econstruction of **A**natomy by **M**apping **O**ntologies*) that appropriately models anatomical dependencies and uses ontology-informed amalgamation of stochastic maps to reconstruct phenotypic evolution at different levels of anatomical hierarchy including entire phenotypes. This approach provides new opportunities for tracking phenotypic radiations and evolution of organismal anatomies.

## INTRODUCTION

Ancestral character state reconstruction has been long used to gain insight into the evolution of individual traits in organisms (Pagel, 1999). However, organismal anatomies (= entire phenotypes) are not merely ensembles of individual traits, rather they are complex systems where traits interact with each other due to anatomical dependencies and developmental constraints. Individual trait approaches substantially simplify the full picture of phenotypic evolution by reducing it to only a single feature at a time, which can potentially hinder the discovery of new evolutionary patterns or even reconstruct logically impossible evolutionary scenarios. Only a handful of studies have been focused on reconstructing evolution of entire phenotypes by treating them as sets of all available traits [e.g., Sauquet et al. (2017); O’leary et al. (2013); Peters et al. (2014)]. Nevertheless, even these studies still employ individual character approaches, which remains the predominant paradigm in comparative phylogenetics due to a lack of methods for modeling an entire phenotype (or its parts) as a single complex character. These limitations thereby prevent researchers from reconstructing entire organismal anatomies. To our knowledge, the only approach that attempts to overcome this problem is the parsimony-based method of Ramírez and Michalik (2014).

In this paper, we propose a new pipeline called *PARAMO* (***P**hylogenetic **A**ncestral **R**econstruction of **A**natomy by **M**apping **O**ntologies*) that takes into account anatomical dependencies and uses stochastic mapping (Huelsenbeck et al., 2003) along with anatomy ontologies to reconstruct the evolution of entire organismal anatomies; this pipeline can be implemented in likelihood or Bayesian frameworks. Our approach treats the entire phenotype or its component body regions as single complex characters and allows exploring and comparing phenotypic evolution at different levels of anatomical hierarchy. These complex characters are constructed by ontology-informed amalgamation of elementary characters (i.e., those coded in character matrix) using stochastic maps. In our approach, characters are linked with the terms from an anatomy ontology, which allows viewing them not just as an ensemble of character state tokens but as entities that have their own biological meaning provided by the ontology.

The goal of this paper is to give the description of *PARAMO* pipeline and *R* (R Core Team, 2018) scripts that can be used to run it. Additionally, we use a Hymenopteran dataset to demonstrate the workflow of the pipeline. At the end of the paper, we discuss biological questions that can be addressed using our method. We believe that reconstructing evolutionary dynamics of entire phenotypes and their major parts opens up new perspectives for comparative morphology and phylogenetics, which, in turn, allows tracking phenotypic radiations across time and phylogeny.

## METHODOLOGICAL BACKGROUND

### The core ingredient: character and character state invariance

At the core of our method lies the property of character and character state invariance that exists in Markov models of discrete trait evolution (Tarasov, 2018, 2019). This property removes the distinction between character and character state, meaning that multiple individual characters can be represented as a single character and vice versa, which makes the two concepts equivalent. In this paper, this invariance property is used to construct larger characters from ensembles of elementary characters, which are constructed through the operation of character amalgamation (see the next section). Character amalgamation is crucial for reconstructing ancestral anatomies because it offers a convenient way to incorporate anatomical dependencies and reconstruct simultaneous evolution of traits at different levels of anatomical hierarchy (= levels of amalgamation).

In present study, we consider three major levels of amalgamation: (1) the level of anatomical dependencies (AD), (2) body regions (BRs) and (3) entire phenotype (EP). The AD level implies that anatomically dependent traits have to be amalgamated into a single character to appropriately model anatomical dependencies (see the Step 2 in the *PARAMO* description). The amalgamation at the BR level implies that all individual characters associated with a particular body region become combined into a single character for comparative analysis. For example, amalgamation of all characters associated with the “head” produces a character that describes evolution of this body region; the same can be done for legs and other BRs of interest. Construction of these characters facilitates comparison of BR evolution across phylogeny. For example, BR amalgamation can be used to address questions of whether different BRs change over the same or different branches on a phylogeny. Amalgamation of all characters in a dataset, produces one gigantic character at the EP level that describes the evolution of an entire anatomy. The character amalgamation has to be performed in a mathematically consistent way, as discussed in the next section.

### Character amalgamation using stochastic maps

In probabilistic models of phylogenetics, a “character” represents a Markov process that sequentially moves from one state to another over time. Discrete characters can be represented as a discrete state Markov process that is defined by a transition rate matrix containing infinitesimal rates of change between states, and an initial vector of probabilities at the root of the phylogenetic tree. Any number of individual characters can be amalgamated into one character through amalgamating their rate matrices (Tarasov, 2019) that defines the joint evolution of the initial characters. In the present paper, we assume that initial characters, if they are not dependent anatomically (see the Step 2 in the *PARAMO* description), are independent entities that have to be independently amalgamated. Suppose there are two characters *C*_1_{with states: *0, 1}* and *C*_2_{with states: *0, 1}* defined by:

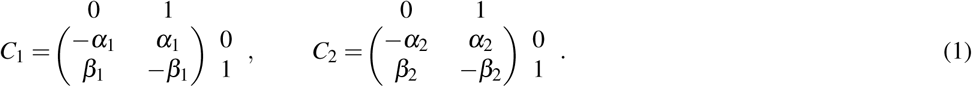

Their independent amalgamation, herein denoted by ⊕ (the Kronecker sum), results in the following character *C*_1,2_ with four states:

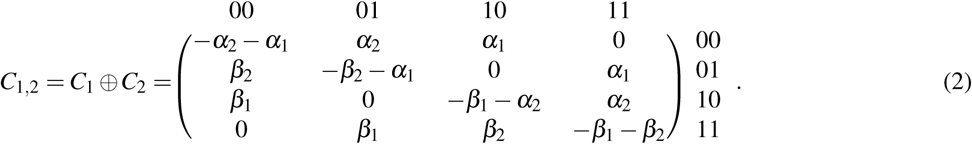

The full formula for character amalgamation can be written as *C*_1_ *⊕ C*_2_ = *C*_1_ *⊗ I_C_*_2_ + *I_C_*_1_ *⊗ C*_2_ where *I_C_*_2_ and *I_C_*_1_ are the identity matrices for *C*_1_ and *C*_2_, and ⊗ denotes the Kronecker product. The amalgamated character *C*_1,2_ {*00,01,10,11*} is constructed by forming its states using the combinations of states from the characters *C*_1_ and *C*_2_. Unfortunately, the rate matrix amalgamation has a shortcoming – the number of its states grows exponentially (as 2^*n*^ for *n* binary characters) resulting in an enormous rate matrix that makes computations infeasible even if it is constructed from a few dozens of initial characters. Here, we propose an approach that bypasses this issue by using stochastic maps.

A stochastic map (*S*) is a phylogenetic tree with an instance of mapped evolutionary history of a character (i.e., state transitions) conditional on data at tips and a Markov model used for ancestral state reconstruction (Huelsenbeck et al., 2003). This tree is divided into segments; each segment corresponds to time spent in a particular state. Thus, stochastic mapping is a function *Sm* that converts realization of a character rate matrix *C* and data to the corresponding stochastic map(s) [i.e., *S* = *Sm*(*C*)]. Both definitions of a character – using rate matrix or stochastic map(s) – are equivalent as they can be converted into each other.

Interestingly, character amalgamation can be performed by only using stochastic maps of the initial characters. Suppose, that *S*_1_ and *S*_2_ are the stochastic maps obtained from the realizations of the characters *C*_1_ and *C*_2_ respectively. The amalgamation of the maps *S*_1_ and *S*_2_ implies construction of a joint stochastic map *S*_1,2_ = *S*_1_ ⊕ *S*_2_ by forming new segments from the combinations of the segments in *S*_1_ and *S*_2_ as shown in (Fig. 1); the map *S*_1,2_ defines the character *C*_1,2_. In other words, the stochastic mapping performed directly on character *C*_1,2_ is identical to the amalgamation of the stochastic maps obtained for *C*_1_ and *C*_2_ separately:

**Figure 1.**
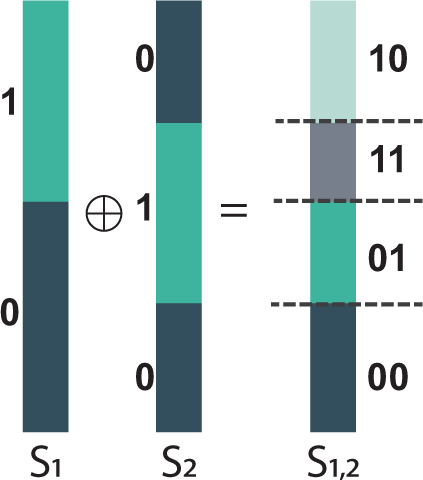
Amalgamation of stochastic maps. Vertical bars are tree branches, their segments are mapped character states. The amalgamation of the stochastic map *S*_1_{0, 1*}* and *S*_2_{0, 1*}* yields the map *S*_1,2_{00, 01, 11, 10*}*.

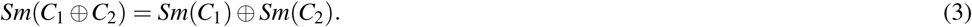

The amalgamation using stochastic maps is computationally cheap as it avoids gigantic rate matrices and can be virtually applied to any number of elementary characters in a dataset. Thus, the invariant property necessary for reconstructing ancestral anatomies can be feasibly maintained. This approach of stochastic map amalgamation is employed in this paper.

### Querying characters using ontologies

Ontologies are graphs that describe relationships (edges) among entities (nodes) from a domain of knowledge under interest. In the present study, we are specifically interested in linking morphological characters with anatomy ontologies to be able to query and retrieve all characters associated with a particular ontology term. Herein, we call this query *”Retrieve all characters”* (*RAC*) and use it to construct character amalgamations for the different levels of the anatomical hierarchy.

In an anatomy ontology, the nodes are the anatomical entities, while edges are their relationships. Linking a character with an ontology, in the context employed here, means assigning a link between ontology term(s) and the character’s ID from a character matrix. Technically, this implies that a character becomes a node in the ontology graph connected with the ontology term(s) its linked to. Two fundamental types of edges – *is a* and *part of* – occur in almost all anatomy ontologies. The relationship *A is a B* indicates that *A* is a subtype of *B*, and the relationship *A part of B* indicates that every instance of *A* is, on the instance level, a part of some instance of *B* (Haendel et al., 2008). Computationally, *RAC* works by taking an input term and traversing the ontology graph using *is a* and *part of* edges to retrieve all characters that are descendant nodes of the input term. In our case, the input is an ontology term that corresponds to a BR or EP. For example, if *RAC* takes the term “head”, it returns all characters associated with this BR (e.g., “shape of eyes”, “length of antennae”, etc.). Thus, ontologies offer a convenient way to automatically query character matrices. The implementation of *RAC* is discussed in the Step 4 section of the pipeline description below.

## DESCRIPTION OF THE PIPELINE

Our pipeline requires three initial pieces of data: a character matrix, a dated phylogeny, and an anatomy ontology. To demonstrate the workflow, we use a modified subset of 9 characters (Table 1) and 87 species from a large-scale phylogeny of Hymenoptera (Sharkey et al., 2012). The character matrix is sketched in Fig. 2A, a detailed description is given in the Supplemental Material. Note, the two pairs of characters in the matrix {*C*_2_*,C*_3_} and {*C*_5_*,C*_6_} are subject to anatomical dependencies (Table 1). For reconstructing character histories, we use the dated phylogeny of Klopfstein et al. (2013), and for linking characters to the ontology, we use the Hymenoptera Anatomy Ontology [HAO, Yoder et al. (2010)]. In this demonstration, we are interested in constructing the amalgamated characters for the AD, BR and EP levels of anatomical hierarchy. At the BR level, three main body regions are considered – “head”, “legs” and “wings” (Fig. 2B).

**Table 1.** Initial characters of Hymenoptera used in demonstration. The symbols *>* and *<* indicate the direction of a hierarchical dependency; the symbol *<>* indicates synchronous dependency (see Step 2 of the pipeline description).

| ID | Character statement | state 0 | state 1 | Dependency |
| --- | --- | --- | --- | --- |
| $C_1$ | Notch on medial margin of eye | absent | present | – |
| $C_2$ | Position of labrum | anterior | posterior | $C_2\{0, 1\} < C_3\{1\}$ |
| $C_3$ | Labrum | absent | present | $C_3\{1\} > C_2\{0, 1\}$ |
| $C_4$ | Forewing costal and radial vein fusion | not fused | fused along their lengths | – |
| $C_5$ | Hind wing subcostal vein, absent | no | yes | $C_5 <> C_6$ |
| $C_6$ | Hind wing subcostal vein, present | yes | no | $C_5 <> C_6$ |
| $C_7$ | Inner posterior mesotibial spur | simple | modified into a calcar | – |
| $C_8$ | Foretibial apical sensillum | present | absent | – |
| $C_9$ | Metatibial apical sensillum | present | absent | – |

**Figure 2.**
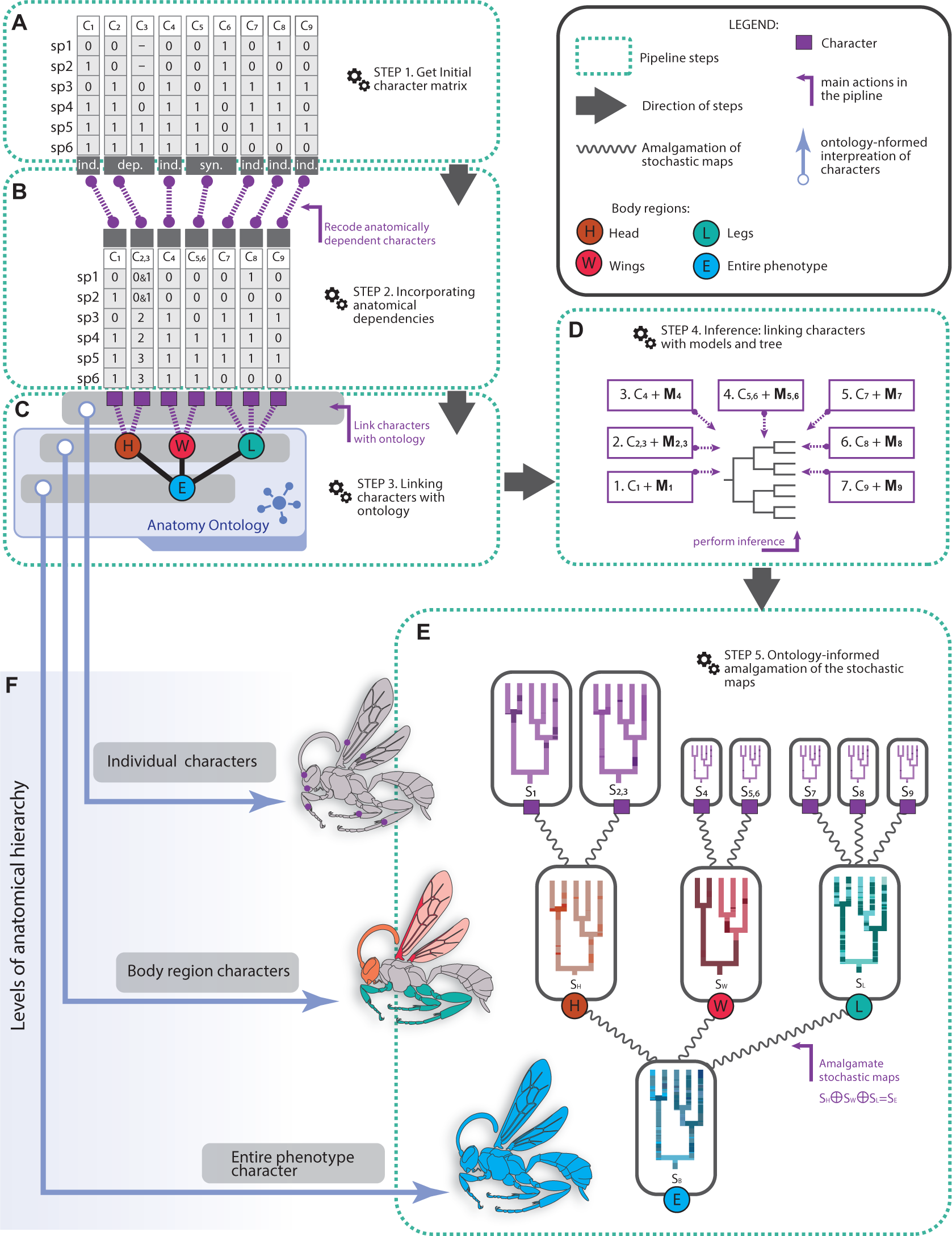
*PARAMO* pipeline. The panels A-E represent the five steps of the pipeline (see the text). E, the size of the stochastic maps *S*_4_-*S*_9_ is reduced for the illustrative purpose. F, three levels of anatomical hierarchy. Abbreviations: *C*-character, *S*-stochastic map, *ind.*-independent character, *dep.* and *syn.*-hierarchically and synchronously dependent characters respectively.

The *PARAMO* pipeline includes five steps as shown in Fig. 2 and described below. In the Supplemental Material, we provide a set of *R* functions that can be used to implement this pipeline in practice and the tutorial (Supp. files *PARAMO pipeline.pdf* or *PARAMO pipeline.Rmd*). The pipeline is also available on *GitHub* https://github.com/sergeitarasov/PARAMO.

### Step 1. Initial character matrix (Fig. 2A)

#### Workflow

The first step requires getting or constructing an initial character matrix that codes a set of characters for a set of species. In our case this matrix is shown in Fig. 2A, and the character report is given in Table

#### Software

Any software for building character matrices can be used at this step, for example, the popular software *Mesquite* (Maddison and Maddison, 2018).

### Step 2. Incorporating anatomical dependencies: constructing amalgamations at the AD level (Fig. 2B)

#### Workflow

The structure of organismal anatomies imposes anatomical dependencies among traits (i.e., the presence of digits is dependent on the presence of limbs). Coding anatomical dependencies has been a subjective procedure because different experts have different views of how to code dependent trait(s) into a character(s). Traditionally, three main coding approaches have been proposed to deal with dependencies: (i) one multistate character, (ii) the presence/absence approach, or (iii) the inapplicable approach. Obviously, AD traits have to be coded appropriately to avoid undesirable bias and incompatible evolutionary states that can negatively affect downstream analyses. The appropriate treatment of anatomical dependencies is discussed in Tarasov (2019) and is followed in the present pipeline. Thus, at this step, we suggest recoding the miscoded AD characters (if there any) from the initial character matrix obtained at Step 1. There two main types of dependencies – hierarchical and synchronous – that appear frequently miscoded in character matrices. Their proper treatment requires use of different coding approaches (Tarasov, 2019).

A *hierarchical dependency* occurs when a hierarchically upstream character controls a downstream one. For example, the state *present*(1) of the character *C*_3_(*Labrum*) controls the presence of both states in *C*_3_(*Position of labrum*), which are inapplicable otherwise (Table 1). The proper way to model this dependency is to use structured Markov models with hidden states that can be constructed by amalgamating the two characters into one as, for instance, shown in Equation 2; this amalgamation results in the character *C*_2,3_ {00, 01, 10, 11} that can be represented using four states as *C*_2,3_ {0, 1, 2, 3}; in *C*_2,3_ the states {0, 1} are hidden and correspond to the observable state *labrum absent*, while the states 2, 3 have the same meaning as those in *C*_2_, respectively. In the character matrix, the hidden states can be scored using polymorphic coding as {0&1} (Fig. 2B). Note, Equation 2 uses the independent amalgamation, other more complex models can be used to model AD characters as well [see Tarasov (2019)].

A *synchronous dependency* usually occurs when a trait is redundantly scored using a binary coding scheme. For example, the characters *C*_5_ and *C*_6_ (Table 1) code the same trait *presence of hind wing subcostal vein* and their character states depend on each other simultaneously: *C*_5_ {0} and {1} occur when *C*_6_ is {1} and {0}, respectively. This synchronous dependency has to be eliminated by combining the two characters into a single character *C*_5,6_ without changing the state pattern (Fig. 2B).

The recoding of dependent characters constructs the amalgamated characters at the AD level. If a character does not display any dependencies then we treat it as correctly amalgamated at the AD level by default. In our demonstrative example, we place the new AD characters in a separate character matrix (Fig. 2B) that is used for the downstream steps in the pipeline.

#### Software

Currently, there is no software that is be capable to automatize the recoding of the AD characters. Obviously, the AD characters can be recode manually using any software for viewing and editing character matrices. If the number of the miscoded characters is large, manual recoding must be taken with caution, as it may result in errors.

### Step 3. Linking characters to an ontology (Fig. 2C)

#### Workflow

In the next step, characters from the previous step must be linked to their respective term(s) from an anatomy ontology. Depending on the scope of a study, the same character might be linked with one or more ontology terms. For the needs of the *PARAMO* approach, the linking requires assigning one or more ontology terms to a respective character as shown in Table 2. The ontology terms have to be selected to best fit a character statement and the scope of a study. In the demonstration here, we are specifically interested in linking the initial characters in a way that facilitates the construction of BD and EP characters.

**Table 2.** Hymenoptera characters linked with HAO terms.

| ID | Character statement | HAO ID | HAO ID name |
| --- | --- | --- | --- |
| C<sub>1</sub> | Notch on medial margin of eye | HAO:0000234 | cranium |
| C<sub>3,2</sub> | Labrum + Position of labrum | HAO:0000639 | mouthparts |
| C<sub>4</sub> | Forewing costal and radial vein fusion | HAO:0000351 | fore wing |
| C<sub>5,6</sub> | Hind wing subcostal vein, present | HAO:0000400 | hind wing |
| C<sub>7</sub> | Inner posterior mesotibial spur | HAO:0001351 | mesotibia |
| C<sub>8</sub> | Foretibial apical sensillum | HAO:0000350 | fore tibia |
| C<sub>9</sub> | Metatibial apical sensillum | HAO:0000631 | metatibia |

For *PARAMO*, character-ontology linking is straightforward methodologically and can be done manually by, for example, constructing a table with the two columns – character ID and an ontology term’s ID (see Table 2). The linking can be facilitated using *R* package *ontoFAST* (https://github.com/sergeitarasov/ontoFAST) that provides a graphical interface for selecting terms by navigating through ontology.

Character-ontology linking, *sensu* this paper, falls into a general area of bioinformatics that focuses on annotating phenotypes with ontologies. The recent developments in this area offer comprehensive methods for constructing detailed annotations of phenotypes and characters (Dahdul et al., 2018; Dececchi et al., 2015; Cui et al., 2016; Balhoff et al., 2013). Such detailed annotations can be constructed in *Phenex* (Balhoff et al., 2010) and used in *PARAMO* as well.

### Step 4. Inference: linking characters with models and tree (Fig. 2D)

#### Workflow

The goal of this step is obtain stochastic maps for the characters amalgamated at the AD level (Step 2). To perform the inference, the characters have to be associated with a dated phylogeny and the respective Markov models of trait evolution. Note, that the anatomically dependent characters require structured Markov models with hidden states (see Step 2); thus these models have to be appropriately assigned to the characters with such dependencies. In our dataset, the only character that requires such model is *C*_2,3_. The stochastic maps can be obtained in likelihood and Bayesian frameworks. The use of the latter is preferable as it provides a convenient way for sampling the stochastic maps from the posterior distribution of character histories, which also incorporates uncertainty.

#### Software

Technically, this step requires creating a data object file(s) for each character that includes the character data, model and tree in an appropriate format that can be read by the software used in inference. In the present tutorial, we use *RevBayes (version 1.0.7)* (Höhna et al., 2016) to perform character inference and generate stochastic maps. The creation of the data files for *RevBayes* is automatized using *R* scripts (Supplemental Material).

### Step 5. Ontology-informed amalgamation of the stochastic maps for AD, BR and EP levels (Fig. 2E)

#### Workflow

Our ultimate goal is to construct the amalgamated characters for the AD, BR and EP levels of the anatomical hierarchy (Fig. 2E). The stochastic maps generated at the previous step are the individual characters of the AD level. The construction of the BR level characters implies the ontology-informed amalgamation of the AD level maps that can be done using a *RAC* query and the characters linked with the ontology in Step 3. In our example, for each of the focal BR terms (”head”, “legs”, wings”), the *RAC* query returns a set of the associated initial characters and their stochastic maps. Next the stochastic maps are used to produce the amalgamated BR characters. The construction of the EP level character is similar to that of BR, but requires amalgamation of all available initial characters.

As soon as the amalgamations are done, this step culminates the pipeline. In the next section, we discuss the use of the amalgamations for addressing various biological questions.

#### Software

The *RAC* query is implemented in *R* using *OntologyIndex* package (Greene et al., 2017) and a set of *PARAMO* functions. The *paramo*() function in the provided *PARAMO* scripts performs the ontology-informed amalgamation of the stochastic maps.

## DISCUSSION

The *PARAMO* pipeline allows users to appropriately incorporate anatomical dependencies and construct characters for phenotypic entities that consist of ensembles of discrete traits. This is achieved through ontology-informed amalgamation of stochastic maps, which allows tracking of evolution at different levels of the anatomical hierarchy – individual characters, body regions (BR) and entire phenotype (EP) (Fig. 2F). Our approach can be applied to a dataset of virtually any size, for example one with hundreds or thousands of characters. Ontology-informed amalgamation at the BR and EP levels represents each entity, usually described by numerous individual characters, as a single multistate character (=single-character representation).

In this paper, we assume that initial characters evolve independently (except those which are anatomically dependent) and hence use independent amalgamation of their stochastic maps. It is known that evolution of morphological characters might be subjected to correlations or more complex scenarios of state changes due to hidden intrinsic or extrinsic factors, which cannot be modeled by the independent amalgamation. In this case, more complex models of trait evolution that incorporate non-anatomical correlations (Pagel, 1994) or hidden states (Beaulieu et al., 2013; Tarasov, 2019) can be also used in *PARAMO*. Similar to modeling anatomical dependencies, these models require appropriately structured rate matrices constructed for a focal set of initial characters at Steps 2 and 4. The application of hidden state models is straightforward in *PARAMO* – a separate hidden model can be assigned to each individual character in the same way the traditional models were used in the provided demonstrative example. In contract, modeling non-anatomical correlation has limitations – it would work well if the focal sets of characters are relatively small that allows using computationally manageable amalgamated rate matrices. However, if the focal sets are large then the amalgamations produce gigantic and computationally intractable matrices thereby precluding the use of the character correlations models. Thus, we emphasize that our method does not resolve the problem of modeling character correlations in morphological dataset to full extent.

Like traditional ancestral state reconstruction, the ancestral states can be also inferred for entire organismal anatomy or its parts through single-character representation. In this case, the states in a large composite BR or EP character correspond to a combination of states from the initial characters that form BR or EP. Note, such integrative reconstruction is similar to the traditional character-by-character reconstruction because *PARAMO* uses independent amalgamation of stochastic maps.

The advantage of *PARAMO* approach is that the single-character representation of BR or EP opens up new avenues for comparing and assessing the dynamics of phenotypic radiations and diversifications. As a response to novel extrinsic or intrinsic factors, a phenotypic radiation may occur by rapidly diversifying an adaptive trait, inherited from a common ancestor, into a diversity of new forms in the ancestor’s descendants. A well-known example of this radiation are Darwin’s finches, which evolved a remarkable diversity of beak shape and functionality. Almost always, such a radiation represents an ensemble of characters that are located on the same or functionally similar body regions. Other body regions (in the same or different species) may also undergo radiations triggered by different factors and coded by their own ensembles of characters. Apparently, these varying ensembles preclude a consistent identification and assessment of phenotypic radiations. In contrast, single-character representation of BRs avoids this problem and may provide insight into the timing, location (clades) and number of phenotypic radiations occurring across a phylogeny. In this respect, each BR character is a stochastic map showing state changes in a tree where the number of changes over a branch (or time interval) reflects the evolutionary rate of the BR character in that branch – in other words, the more changes the faster the rate. The per-branch rate estimates can be used to determine rate shifts in the BR character and identify the timing of an evolutionary radiation. The same approach applied to a set of BRs can be used to map different phenotypic radiations onto the phylogeny. Obviously, any organism has many BRs that are hierarchically structured due to the nature of the anatomy. Ontology-informed amalgamation can generate characters for all potential BRs, thereby allowing to study phenotypic radiations hierarchically [see Slater and Friscia (2019)]. Additionally, the single-character representation of EP can be used to identify rate shifts in the entire organismal anatomy and address questions on tempo and mode of its evolution. Thus, *PARAMO* can be used to disentangle evolution of phenotypes across tree and body regions. Unfortunately, by far, identification of rate shifts using amalgamated stochastic maps requires new statistical methods. Their development is beyond the focus of present study but we anticipate their emergence in near future.

The decades of systematists’ effort have generated thousands of datasets (see *Phenoscape* (Mabee et al., 2012) and *Morphobank* (O’Leary and Kaufman, 2011)), that score morphological characters for numerous clades across the Tree of Life. Frequently, these morphological data become forgotten shortly after publishing a phylogenetic tree they were used to construct. The *PARAMO* approach and anticipated further development in this area provide a new dimension for analyzing these data, which, as we believe, will aid understanding of how phenotypes evolve.

## ACKNOWLEDGMENTS

We are grateful to Brian O’Meara (University of Tennessee) for helping to obtain the dated tree of Hymenoptera.

## REFERENCES

Balhoff, J. P., Dahdul, W. M., Kothari, C. R., Lapp, H., Lundberg, J. G., Mabee, P., Midford, P. E., Westerfield, M., and Vision, T. J. (2010). Phenex: ontological annotation of phenotypic diversity. PLoS One, 5(5):e10500.

Balhoff, J. P., Mikó, I., Yoder, M. J., Mullins, P. L., and Deans, A. R. (2013). A semantic model for species description applied to the ensign wasps (hymenoptera: Evaniidae) of new caledonia. Systematic Biology, 62(5):639–659.

Beaulieu, J. M., O’meara, B. C., and Donoghue, M. J. (2013). Identifying hidden rate changes in the evolution of a binary morphological character: the evolution of plant habit in campanulid angiosperms. Systematic biology, 62(5):725–737.

Cui, H., Xu, D., Chong, S. S., Ramirez, M., Rodenhausen, T., Macklin, J. A., Ludäscher, B., Morris, R. A., Soto, E. M., and Koch, N. M. (2016). Introducing explorer of taxon concepts with a case study on spider measurement matrix building. BMC bioinformatics, 17(1):471.

Dahdul, W., Manda, P., Cui, H., Balhoff, J. P., Dececchi, A., Ibrahim, N., Lapp, H., Vision, T., and Mabee, P. M. (2018). Annotation of phenotypes using ontologies: a gold standard for the training and evaluation of natural language processing systems. bioRxiv, page 322156.

Dececchi, T. A., Balhoff, J. P., Lapp, H., and Mabee, P. M. (2015). Toward synthesizing our knowledge of morphology: Using ontologies and machine reasoning to extract presence/absence evolutionary phenotypes across studies. Systematic biology, 64(6):936–952.

Greene, D., Richardson, S., and Turro, E. (2017). ontologyx: a suite of r packages for working with ontological data. Bioinformatics, 33(7):1104–1106.

Haendel, M. A., Neuhaus, F., Osumi-Sutherland, D., Mabee, P. M., Mejino, J. L., Mungall, C. J., and Smith, B. (2008). Caro–the common anatomy reference ontology. In Anatomy Ontologies for Bioinformatics, pages 327–349. Springer.

Höhna, S., Landis, M. J., Heath, T. A., Boussau, B., Lartillot, N., Moore, B. R., Huelsenbeck, J. P., and Ronquist, F. (2016). Revbayes: Bayesian phylogenetic inference using graphical models and an interactive model-specification language. Systematic Biology, 65(4):726–736.

Huelsenbeck, J. P., Nielsen, R., and Bollback, J. P. (2003). Stochastic mapping of morphological characters. Systematic Biology, 52(2):131–158.

Klopfstein, S., Vilhelmsen, L., Heraty, J. M., Sharkey, M., and Ronquist, F. (2013). The hymenopteran tree of life: evidence from protein-coding genes and objectively aligned ribosomal data. PLoS One, 8(8):e69344.

Mabee, P., Balhoff, J. P., Dahdul, W. M., Lapp, H., Midford, P. E., Vision, T. J., and Westerfield, M. (2012). 500,000 fish phenotypes: The new informatics landscape for evolutionary and developmental biology of the vertebrate skeleton. Journal of Applied Ichthyology, 28(3):300–305.

Maddison, W. and Maddison, D. (2018). Mesquite: a modular system for evolutionary analysis. version 3.40. available from: http.mesquiteproject.org (accessed 15 January 2018).

O’leary, M. A., Bloch, J. I., Flynn, J. J., Gaudin, T. J., Giallombardo, A., Giannini, N. P., Goldberg, S. L., Kraatz, B. P., Luo, Z.-X., Meng, J., et al. (2013). The placental mammal ancestor and the post–k-pg radiation of placentals. Science, 339(6120):662–667.

O’Leary, M. A. and Kaufman, S. (2011). Morphobank: phylophenomics in the “cloud”. Cladistics, 27(5):529–537.

Pagel, M. (1994). Detecting correlated evolution on phylogenies: a general method for the comparative analysis of discrete characters. Proc. R. Soc. Lond. B, 255(1342):37–45.

Pagel, M. (1999). The maximum likelihood approach to reconstructing ancestral character states of discrete characters on phylogenies. Systematic biology, 48(3):612–622.

Peters, R. S., Meusemann, K., Petersen, M., Mayer, C., Wilbrandt, J., Ziesmann, T., Donath, A., Kjer, K. M., Aspöck, U., Aspöck, H., et al. (2014). The evolutionary history of holometabolous insects in-ferred from transcriptome-based phylogeny and comprehensive morphological data. BMC evolutionary biology, 14(1):52.

R Core Team (2018). R: A Language and Environment for Statistical Computing. R Foundation for Statistical Computing, Vienna, Austria.

Ramírez, M. J. and Michalik, P. (2014). Calculating structural complexity in phylogenies using ancestral ontologies. Cladistics, 30(6):635–649.

Sauquet, H., von Balthazar, M., Magallón, S., Doyle, J. A., Endress, P. K., Bailes, E. J., de Morais, E. B., Bull-Hereñu, K., Carrive, L., Chartier, M., et al. (2017). The ancestral flower of angiosperms and its early diversification. Nature communications, 8:16047.

Sharkey, M. J., Carpenter, J. M., Vilhelmsen, L., Heraty, J., Liljeblad, J., Dowling, A. P., Schulmeister, S., Murray, D., Deans, A. R., Ronquist, F., et al. (2012). Phylogenetic relationships among superfamilies of hymenoptera. Cladistics, 28(1):80–112.

Slater, G. J. and Friscia, A. R. (2019). Hierarchy in adaptive radiation: a case study using the carnivora (mammalia). Evolution.

Tarasov, S. (2018). The invariant nature of a morphological character and character state: Insights from gene regulatory networks. bioRxiv, page 420471.

Tarasov, S. (2019). Integration of anatomy ontologies and evo-devo using structured Markov models suggests a new framework for modeling discrete phenotypic traits. Systematic biology.

Yoder, M. J., Miko, I., Seltmann, K. C., Bertone, M. A., and Deans, A. R. (2010). A gross anatomy ontology for hymenoptera. PloS one, 5(12):e15991.

